# Genomic epidemiology of *Corynebacterium diphtheriae* in New Caledonia

**DOI:** 10.1101/2022.10.23.512725

**Authors:** Eve Tessier, Melanie Hennart, Edgar Badell, Virginie Passet, Julie Toubiana, Antoine Biron, Ann-Claire Gourinat, Audrey Merlet, Julien Colot, Sylvain Brisse

**Author notes:** **Correspondence: Sylvain Brisse:** Institut Pasteur, Biodiversity and Epidemiology of Bacterial Pathogens, 28 rue du Docteur Roux, F-75724 Paris, France.

## Abstract

**Objectives:** An increasing number of isolations of *Corynebacterium diphtheriae* has been observed in recent years in the archipelago of New Caledonia. We aimed to analyze the clinical and microbiological features of samples with *C. diphtheriae*.

**Methods:** All *C. diphtheriae* isolates identified in New Caledonia from May 2015 to May 2019 were included. For each case, a retrospective consultation of the patient files was conducted. Antimicrobial susceptibility phenotypes, *tox* gene and diphtheria toxin expression, biovar and the genomic sequence were determined. Core genome multilocus sequence typing (cgMLST), 7-gene MLST and search of genes of interest were performed from genomic assemblies.

**Results:** 58 isolates were included, with a median age of patients of 28 years (range: 9 days to 78 years). Cutaneous origin accounted for 51 of 58 (87.9%) isolates, and *C. diphtheriae* was associated with *Staphylococcus aureus* and/or *Streptococcus pyogenes* in three quarters of cases. Half of cases came either from the main city Noumea (24%, 14/58) or from the sparsely populated island of Lifou (26%, 15/58). Six tox-positive isolates were identified, associated with recent travel to Vanuatu; 5 of these cases were linked and cgMLST confirmed recent transmission. Two cases of endocarditis in young female patients with a history of rheumatic fever involved tox-negative isolates. The 58 isolates were mostly susceptible to commonly used antibiotics. In particular, no isolate was resistant to the first-line molecules amoxicillin or erythromycin. Resistance to tetracycline was found in a genomic cluster of 17 (29%) isolates, 16 of which carried the *tetO* gene. There were 13 cgMLST sublineages, most of which were also observed in the neighboring country Australia.

**Conclusions:** Cutaneous infections may harbor non-toxigenic *C. diphtheriae* isolates, which circulate largely silently in non-specific wounds. The possible introduction of tox-positive strains from a neighboring island illustrates that diphtheria surveillance should be maintained in New Caledonia, and that immunization in neighboring islands must be improved. Genomic sequencing uncovers how genotypes circulate locally and across neighboring countries.

## Introduction

Once a major cause of child mortality, diphtheria has been largely controlled following mass vaccination (1). However, 4,000 to 8,000 cases are reported each year worldwide (1, 2). Large recent outbreaks have been linked to insufficient local vaccination coverage rate following economic crises, political unrest and population displacements (3–5). Diphtheria is an infection caused mainly by *C. diphtheriae,* which is transmitted among humans. The disease has two main forms, pseudomembranous tonsillitis and cutaneous diphtheria. Other clinical presentations include systemic syndromes caused by the action of diphtheria toxin, endocarditis and secondary localizations due to hematogenous dissemination of non-toxigenic isolates (6, 7). Only some *C. diphtheriae* strains carry the *tox* gene, which codes for the diphtheria toxin. Although the bacterium typically transmits through droplets in respiratory infections, transmission can also occur through contact with infected or colonized skin lesions. Cutaneous diphtheria has become more frequent than the classical respiratory form in developed countries, and probably plays an important role during inter-epidemic periods in the tropics, where sporadic cases occur frequently.

The spatial distribution, local spread and global dissemination of C. *diphtheriae* strains are largely undocumented. Genomic sequencing used in the context of epidemiological surveillance can help decipher the links between isolates and infer patterns of spread. So far, apart from outbreak situations (8, 9, 5), few genomic epidemiology studies of diphtheria have been performed (10–14).

New Caledonia, an overseas collectivity of France, is located in the South Pacific region; it is an island with tropical weather and a population of about 270, 000 people (15). This island system makes it an interesting case study to describe the diversity of *C. diphtheriae* isolates and their spread within the island and into or from it. Links with cases of diphtheria in neighboring countries such as Indonesia (16, 17) and Australia (18, 19) are unknown. Frequent travel between New Caledonia and mainland France may be accompanied by *C. diphtheriae* dissemination (12).

The aim of this study was to describe clinical, microbiological and genotypic characteristics of *C. diphtheriae* strains isolated in New Caledonia during four years. Clinical presentations, geographical localization and microbiological features of 58 cases of infection or carriage of *C. diphtheriae* were reported, and a genomic comparison of isolates from New Caledonia and other world regions was performed.

## Material and methods

### Inclusion and exclusion criteria

Local legislation obliges all laboratories of New Caledonia to report and send any isolated *C. diphtheriae* to the Centre Hospitalier Territorial (CHT) microbiology reference laboratory. Cases were defined as an infection or carrier with an isolate identified as *C. diphtheriae,* by the Matrix-Assisted Laser Desorption Ionization Time of Flight Mass Spectrometry (MALDI-TOF MS) automated system (Microflex Bruker). We investigated all episodes with *C. diphtheriae* isolated in culture (n=57 from New Caledonia, plus one from Vanuatu; see below) over a 4 years period (from May 2015 to May 2019). The included cases were issued from clinical samples taken at the main hospital (CHT, n=36) or from *C. diphtheriae* isolates sent to the reference Laboratory by other private and public laboratories located in New Caledonia (in this study, by two contributing laboratories; n = 21).

No clinical criteria for inclusion or exclusion were retained. In most cases (53/58), bacteriological examination was triggered by symptomatic manifestations. All initial cases (including the tox-negative ones, as investigations were triggered before the *tox* PCR result) were investigated after notification. In two investigations, further *C. diphtheriae* isolates were found: in investigation group 1 **(Table 1**), one tox-negative isolate (FRC0493) was found in a contact (case 29) of a tox-negative index case (case 28 in **Table 1**). In investigation group 2, four tox-positive isolates from contacts of an index case (case 60 in **Table 1**), detected by pharyngeal or skin lesion swabbing, and one tox-negative isolate (case 86, a contact in Vanuatu), were found. The latter case was included in the study, despite being from Vanuatu. Hence, 58 cases were included in total.

### Patient record and clinical data

For each patient at CHT, a retrospective investigation in the medical records was carried out and relevant information was compiled in **Table 1** (see also **Table S1**). For patients not recorded in the care software or whose diphtheria episode was not registered, a manual search in the records was carried out. For cases detected by laboratories outside the CHT, the prescribers were contacted in order to access the medical records. As the two dispensaries on the island of Lifou were the prescriber for a large number of cases, an on-site visit was organized and the missing data were recovered by consulting their records. Finally, the data collected by the Direction of Sanitary and Social Affairs (DASS) during each epidemiological investigation around a case, completed eventually the initial data.

### Antimicrobial susceptibility testing

Antimicrobial susceptibility testing (AST) analyses were performed at the CHT laboratory after subculture of the strain on blood agar and verification of the identification by MALDI-TOF (Bruker). A 0.5 McFarland suspension was made in a saline solution (physiological serum), following the recommendations of the Comité de l’Antibiogramme de la Société Française de Microbiologie (CASFM), 2019 version. To determine the zone diameter for erythromycin and azithromycin, a 15 μg disk was used; for penicillin G, a 10 units disk content was used in addition to the disk with 1/10 dilution recommended by the CASFM 2013 version. The agar plates (Mueller-Hinton MHF, BioMérieux, Marcy l’Etoile, France) were inoculated by swabbing. After deposition of antibiotic paper discs (BioRad) and E-test strips (BioMérieux, Marcy l’Etoile, France), the plates were incubated 18 to 24 h at 35±2°C, under an atmosphere enriched with 5% CO_2_. The reading was performed manually, by the same operator, using a caliper. The names of antimicrobial agents, dosages, incubation conditions and interpretation thresholds are given in **Table S2.**

### Diphtheria toxin gene presence (*tox* positivity) and toxin production (toxigenicity)

All strains were tested for the presence of the *tox* gene at the CHT reference laboratory by real-time PCR, using the primers and probe from Schuhegger et *al*. (20): Cdiph-RTDT_Fw primer: 5’-TTA-TCA-AAA-GGT-TCG-GTG-ATG-GTG-3’; Cdiph-RTDT_rev2 primer: 5’-AAT-CTC-AAG-TTC-TAC-GCT-TAA-C-3’; and Cdiph-RTDT_So probe: 6-FAM-5’CGC-GTG-TAG-TGC-TCA-GCC-TTC-CCT-3’-TAMRA. Amplification conditions were modified by using the Roche LightCycler 2.0 (LC 2.0) thermal cycler, with a denaturation step (10 min, 95°C) and 45 cycles of amplification: denaturation for 15 seconds at 95°C and hybridization/elongation for 30 seconds at 60°C. For each assay, positive and negative controls were used. DNA extraction were carried out using MagNA Pure LC total Nucleic acid isolation kit on the automated MagNA Pure LC2.0 platform (Roche) from two to five colonies picked up from fresh cultured plates.

All isolates carrying the *tox* gene were tested at the national reference center at Institut Pasteur for toxigenicity, using the modified Elek immuno-precipitation test (21).

### Biovar determination

Strains were characterized biochemically for pyrazinamidase, urease, and nitrate reductase and for utilization of maltose and trehalose using API Coryne strips (BioMérieux, Marcy l’Etoile, France) and the Rosco Diagnostica reagents (Eurobio, Les Ulis, France). The Hiss serum water test was used for glycogen fermentation. The biovar of isolates was determined based on the combination of nitrate reductase (positive in biovars Mitis and Gravis, negative in biovar Belfanti) and glycogen fermentation (positive in biovar Gravis only).

### Genome sequencing and phylogenetic analyses

The DNA preparation, sequencing and the *de novo* assembly follows the same protocol as the previous studies (12, 22). The taxonomy of the isolates was confirmed by a MASH (v2.2) genomic distance lower than 0.05 with *C. diphtheriae* type strain NCTC11397^T^ or *C. belfantii* type strain FRC0043^T^.

We built a core genome multiple sequence alignment (cg-MSA) from the assembled genome sequences. For this, the genome sequences were annotated using PROKKA v1.14.5 (23) with default parameters, resulting in GFF files. Panaroo v1.2.3 (24) was used to define proteincoding gene clusters, with a threshold set at 80% amino acid identity, then the proteincoding gene sequences of each locus were aligned using MAFFT v7.467 (25) with default parameters. Core genes were defined as being present in 95% of genomes and were concatenated into a cg-MSA by Panaroo. IQtree version 2 (26) with best-fit model GTR+F+R7 was used to build a tree.

We used AMRFinderPlus v3.10.20 to identify known resistance genes in *C. diphtheriae* genomic sequences, based on the December 21^st^, 2021 database and using the following parameters to define presence of a gene: identity >□90% and length coverage >□50%. We used BLASTN (identity >□90% and coverage >□50%) to search for the presence of the *tox* gene, *spuA* gene (associated with biovar Gravis) and *narG* (associated with biovar Belfantii).

MLST and cgMLST genotypes were defined using the Institut Pasteur *C. diphtheriae* and *C. ulcerans* database at https://bigsdb.pasteur.fr/diphtheria. We used a previously described cgMLST scheme, database and associated classification schemes (22) for genomic clusters (25 mismatch threshold) and sublineages (500 mismatches). The isolates provenance data and genome sequences were imported into an isolates database (https://bigsdb.pasteur.fr/cgi-bin/bigsdb/bigsdb.pl?db=pubmlst_diphtheria_isolates&page=query). A publicly available BIGSdb project (i.e., a browsable list of isolates) entitled ‘New Caledonia 2015-2019 study’, corresponding to the present dataset, was created to facilitate retrieval and analysis of the data.

### Data statement

The genomic sequences generated in this study were deposited in the ENA archive and are available from the INSDC databases (ENA/NCBI/DDBJ) under accession numbers: see **Table S3** (*in process for some of them*).

### Ethical statement

This work was approved by the CHT ethics committee (Avis 2022-001).

## Results

### Demographic and clinical data

The demographic and clinical characteristics of the cases are presented in **Table** *1;* 58 cases were included, with a sex ratio of 1.07 (30 men and 28 women). The median age of subjects was 28 years (range, 9 days – 78 years). The geographic distribution of cases **(Figure 1**) was uneven; the two districts with most cases were Nouméa (14 cases, 24.1%) and Lifou island (15 cases, 25.9%).

**Figure 1.**
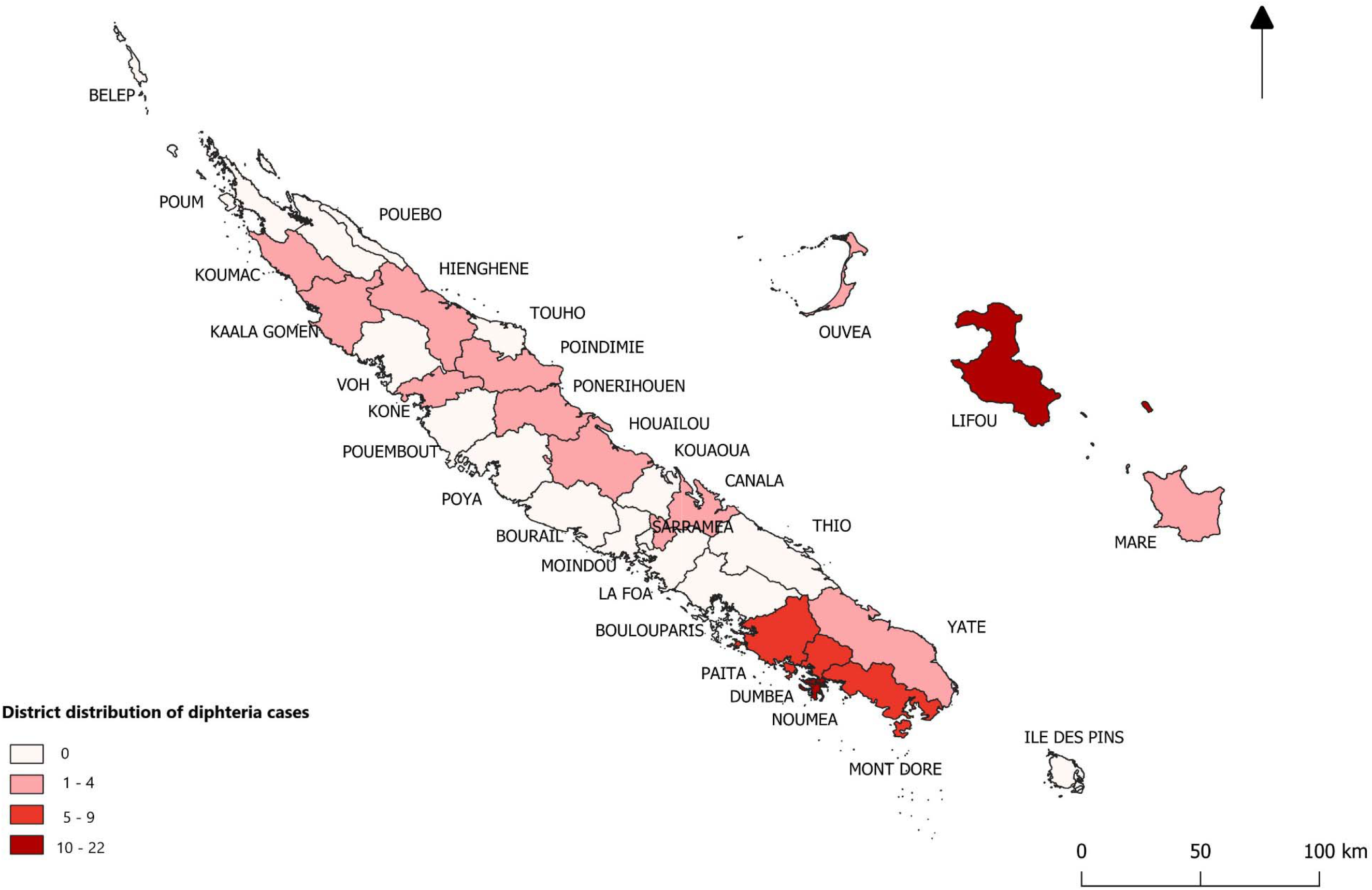
Geographic distribution of *Corynebacterium diphtheriae* cases in New Caledonia. The number of patients whose living place is in each district, is indicated as a heatmap (see key). The arrow indicates the direction of the North.

A large majority of cases (51/58, 87.9%) corresponded to polymicrobial wounds or abscesses, in association with S. *aureus* (35/58, 60.3%) and/or S. *pyogenes* (31/58, 53.4%) **(Table S1).** For the remaining patients, *C. diphtheriae* was found in 1 urine, 2 throat samples, 1 implantable site, and 3 blood cultures; the three latter isolates were found in pure culture. The urine sample was collected because of fever of unknown origin (but without any urinary functional sign). Positive blood cultures were associated with an infective endocarditis in 2 cases, while the third case with positive blood culture was probably a cutaneous contamination. All carriers of tox-positive *C. diphtheriae* were properly vaccinated.

### Microbiological characteristics

The search of the *tox* gene by PCR showed that most isolates (52/58, 89.7%) were tox-negative. The six tox-positive isolates corresponded to cases imported from Vanuatu. Elek’s toxigenicity test showed that all of these isolates were toxigenic, *i.e.,* produced the toxin **(Table 1)**.

Antimicrobial susceptibility testing **(Table S1)** showed that all isolates were susceptible to amoxicillin, ciprofloxacin, rifampicin, cotrimoxazole, erythromycin, azithromycin, vancomycin and clindamycin (except one). For tetracycline, 17 resistant isolates were observed. Genomic analysis of antimicrobial resistance genes showed the presence of *tetO* gene in the 16 isolates of sublineage SL416 **(Table S1).** All isolates with *tetO* were resistant to tetracycline, and all isolates without *tetO* gene were susceptible to tetracycline except one (FRC0758); for this isolate, no explanation was found for its tetracycline resistance phenotype.

The interpretations for penicillin G depended on the reference system used: All isolates were resistant based on the 2019 CASFM reference system **(Table S2),** but using the 2013 CASFM guidelines, only 5 isolates were classified as intermediate, the others being classified as susceptible **(Table S1).**

The *pbp2m* gene was observed in one isolate (FRC0534), with decreased susceptibility to penicillin (CMI: 0.38 g/L). This gene is typically found on a small genetic unit, sometimes associated with *ermX,* in several plasmidic or chromosomal contexts, but is rare in *C. diphtheriae* populations (12, 27). Here, it was not associated with *ermX,* but with a nearby relaxase gene as described previously (12). No macrolide resistance gene was found among the studied isolates.

The genomic analyzes confirmed the MALDI-TOF species identification as *C. diphtheriae* for 57 isolates, whereas one tox-negative isolate (FRC0723) was identified as belonging to the closely related species *C. belfantii*. Biotyping showed that two isolates were of biovar Belfanti (including the *C. belfantii* isolate). Among the remaining isolates, 46 were of biovar Gravis and 10 of biovar Mitis **(Table S1).**

Biovars Mitis and Gravis are distinguished by the ability to utilize glycogen. The *spuA* gene was reported as being associated with biovar Gravis isolates (28, 12). We confirmed a strong association between the presence of *spuA* and the Gravis phenotype: almost all (42/46; 91.3%) isolates having the *spuA* gene were of Gravis phenotype **(Figure 2).** No molecular explanation was found for the ability to utilize glycogen of the four other Gravis isolates, and we noted that in some of them, the *spuA* gene was truncated **(Table S1).**

**Figure 2.**
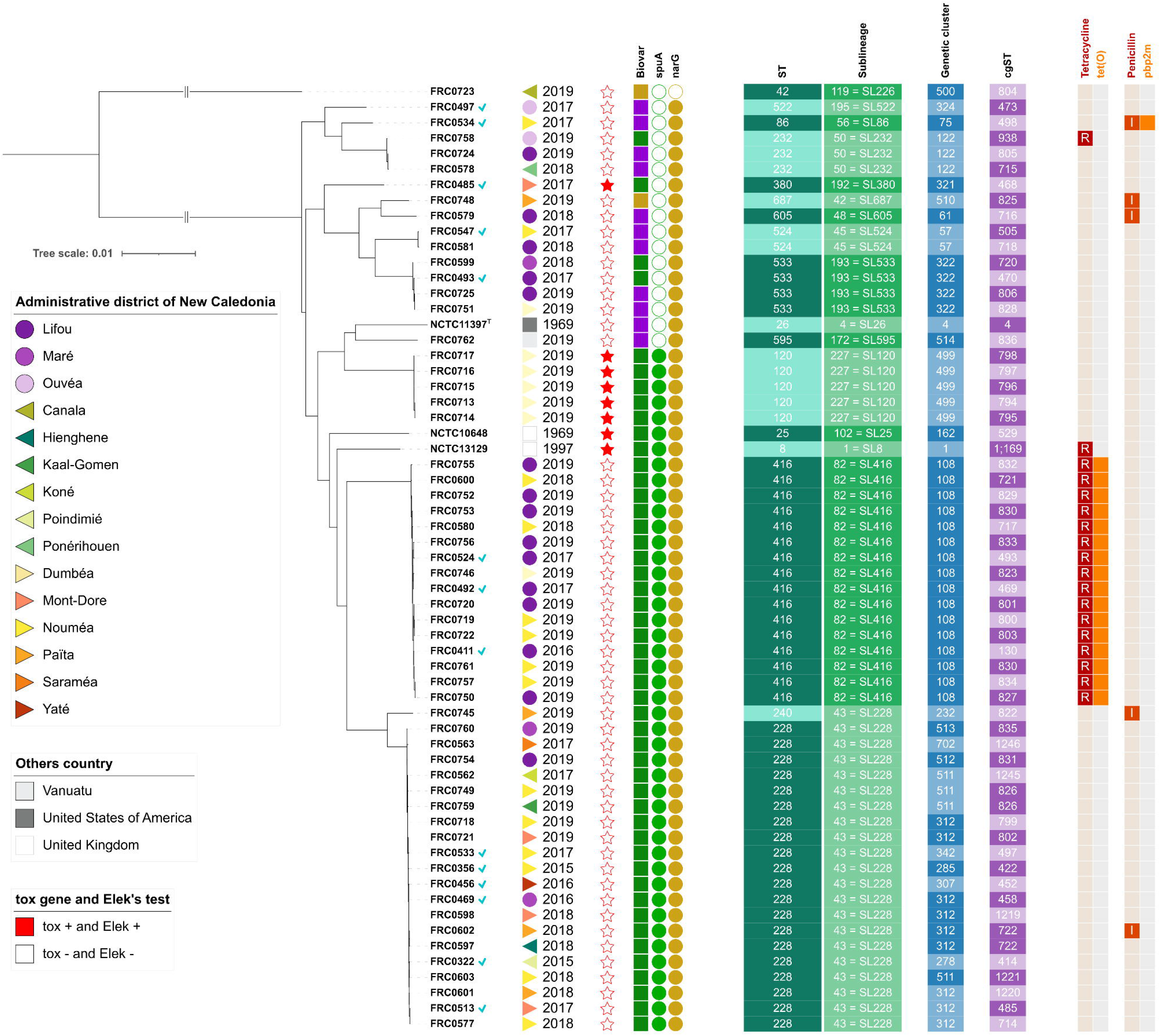
Diversity of *Corynebacterium diphtheriae* isolates from New Caledonia. The first column after the isolate identifier represents the locations of origin of the isolates (see color key). After the year of isolation column, the tox-negative isolates are represented by an empty star, and tox-positive ones by a full red star. The biovar is represented next: green for Gravis, purple for Mitis, brown for Belfanti. The following columns represent the presence of genes *spuA* (likely involved in glycogen metabolism; positive in biovar Gravis) and *narG* (involved in nitrate metabolism; positive in biovars Gravis and Mitis). On the four next columns, alternating dark and light colors indicate each sequence type (ST), sublineage (SL), genetic cluster (GC) or cgST change. Sublineages are given as: internal number = alias number (inherited, where possible, from 7-gene MLST numbers). The next column represents the tetracycline resistance phenotype: red for resistant (with R letter), grey for susceptible. The presence of the *tetO* gene is symbolized in orange on the next column, its absence in grey. The last two columns indicate in the same way, the penicillin phenotype and presence of gene pbp2m (I: intermediate).

The nitrate reductase activity differentiates Mitis and Gravis isolates from biovar Belfanti isolates, which are nitrate-negative. As expected (29, 30), the *narG* gene (coding for nitrate reductase) was absent in the FRC0723 *C. belfantii* strain. However, no genomic explanation was found for the nitrate-negative phenotype of *C. diphtheriae* biovar Belfanti isolate (FRC0748).

### Genomic epidemiology of isolates

*C. diphtheriae* isolates can be classified into 7-gene MLST sequence types (31) and into phylogenetic sublineages, which represent deep phylogenetic subdivisions of the population structure of this species that are highly concordant with ST classifications (22). The 7-gene MLST sequence type (ST) and cgMLST classifications into sublineages (SL; maximum of 500 mismatches out of 1305 loci) were defined for the 58 isolates **(Table 1; Figure 2).** Based on MLST, there were 14 distinct STs, whereas cgMLST grouped the isolates into 13 different sublineages (SL). There was total agreement between ST and SL classifications, except that one isolate of sublineage SL228 had ST240 instead of ST228 **(Table** 1). Sublineage SL228 was represented by 21 isolates (36.2% of the total) and sublineage SL416 was represented by 16 (27.6%) isolates **(Table 1)**. Hence, these two predominant sublineages represented more than half of the isolates circulating in New Caledonia. Isolates of these two sublineages were all of biovar Gravis. Whereas SL228 was isolated in 10 administrative districts, SL416 isolates were all recovered from Nouméa and Lifou except one **(Table *1;* Figure 2).**

Genomic clusters (GC) are much narrower genetic subdivisions of sublineages that have been defined as groups of isolates that have, among themselves, genetic distances (maximum of 25 cgMLST mismatches) compatible with outbreaks or recent transmission (22). Looking at this genotyping classification level, there were 22 distinct GCs **(Table 1; Figure 2).** Two of these were recovered more than 8 times: GC108 (16 isolates) and GC312 (9 isolates), and they belonged to each of the predominant sublineages (SL416 and SL228), respectively. However, whereas SL228 comprised 9 other GCs, SL416 comprised only GC108, showing that SL416 is genetically very homogeneous, and may correspond to relatively recent transmission. In contrast, SL228 is genetically heterogeneous, in line with much wider geographic distribution across the island **(Table 1)**. There was no documented epidemiological link between isolates in these genomic clusters. Hence, they may correspond to cryptic transmission that was revealed here by genomic analyses.

Genetic cluster GC499 corresponded to tox-positive isolates from an index case and isolates from contacts that were screened during investigations around the index case (Investigation group 2 in **Table 1**). In this investigation, samples were collected in the same family in New Caledonia, of which some members had a recent history of travel to Vanuatu. Further comparisons of cgMLST allelic profiles within CG499 showed a maximum of 2 mismatches among the five isolates, strongly concordant with recent transmission of a single strain. Another contact isolate (case 86, isolate FRC0762) collected in Vanuatu during this investigation was in fact not related to the others based on genomic analyses: the isolate was tox-negative and belonged to a distinct sublineage **(Table 1**).

Another epidemiological investigation was triggered by a tox-negative isolate from a case on Lifou island. Upon screening of contacts, one isolate was found (see ‘Investigation group 1’ in **Table 1**). Unexpectedly, the isolates from the index and the contact were in fact genetically unrelated, as they belong to different STs and sublineages (and of course, genetic clusters). The index isolate (FRC0492) belonged to the predominant genomic cluster circulating in Lifou island (GC108).

Genotypic data of the isolates from New Caledonia were compared with those of 876 global *C. diphtheriae* isolates available in public genome data repositories. Of the 13 sublineages from New Caledonia, 8 were also observed elsewhere **(Table S3).** Seven of these were observed in Australia, 2 in mainland France, and 3 in other European countries. Australian genomes represent only 66 (4.5%) of comparative genomes, and are therefore clearly overrepresented in sublineages shared with New Caledonia. Remarkably, two genomic clusters represented in New Caledonia, GC57 and GC61, were also reported 2012-2015 in Australia **(Table S3),** indicative of relatively recent transmission between the two islands.

## Discussion

We provide clinical and microbiological data on a series of 58 infection or carriage cases of *C. diphtheriae* isolates in New Caledonia. Nearly 90% of the samples corresponded to skin infections, and *C. diphtheriae* was found in association with the pathogens S. *aureus* or S. *pyogenes* in over three quarters of these cases. Furthermore, the vast majority of *C. diphtheriae* isolates were non-toxigenic, which puts the pathogenicity of C. *diphtheriae* in skin infections into perspective. This study suggests that *C. diphtheriae* could be pathogenic itself or can colonize a pre-existing infection caused by another bacterium. Co-infection could contribute to an increase of severity, compared to the same infection without *C. diphtheriae*. The classic clinical presentation is an ulcer covered with a pseudomembrane (32, 33). The pathogenicity of *C. diphtheriae* strains in this kind of context is not proved and the treatment should cover other pathogenic bacteria. The only two isolates from throat swabs were obtained from asymptomatic patients, in the scope of a confirmed case investigation. The strain of urinary origin was probably a contamination during the urine collection.

*C. diphtheriae* endocarditis is well described (34, 35). This infection frequently occurs in patients with predispositions, such as pre-existing heart disease, prosthetic valves and intravenous drug use (35). Cases of *C. diphtheriae* endocarditis have been described in subjects who are generally younger than described for other pathogens. Endocarditis strains may or may not carry the diphtheria toxin gene, but cases of tox-negative endocarditis have predominated since high global vaccination coverage has been achieved (34, 35). Here, both cases of infective endocarditis were in young women with a history of rheumatic fever, a common condition in New Caledonia (36). People belonging to Melanesian and Polynesian populations, which are more affected by rheumatic fever than those of the European population, are at greater risk of developing infective endocarditis (36). Here, we showed that *C. diphtheriae* is a contributor; of note, six other cases of *C. diphtheriae* endocarditis were previously reported in New Caledonia, five of them between 2005 and 2011, and one in 2021 (CHT laboratory data, unpublished), for a total of 8 cases, and with a history of acute rheumatoid fever for 6 patients.

A large majority of isolates were tox-negative strains. The only tox-positive isolates were from patients returning from Vanuatu, and were all toxigenic based on Elek’s test. Toxigenic isolates can cause severe respiratory or skin infections, and the action of the toxin can lead to cardiac dysfunction and neurological damage. In the present cases, no toxinic symptoms were observed.

Vaccination coverage among children is insufficient in Vanuatu, with only 62% of children being vaccinated in 2021 (37). Improving current vaccination coverage in Vanuatu would seem important, and a study of the epidemiological situation of diphtheria in this setting could guide control measures. Although carriers of tox-positive *C. diphtheriae* in New Caledonia were properly vaccinated, the immune response is directed against the toxin, and although it is effective against toxinic symptoms, it is not considered effective at preventing colonization.

The isolates were all susceptible to amoxicillin, and erythromycin, two first-line molecules for diphtheria treatment. For penicillin G, the French CA-SFM recommendations changed dramatically in 2014, in line with the European Committee on Antimicrobial Susceptibility Testing (EUCAST). These changes cover susceptibility testing protocols, disk loads used and interpretation thresholds. The new 1 IU disc load and 29 mm threshold classifies *de facto* all isolates as “resistant”. Conversely, the 2013 version classified almost all isolates in this study as “susceptible” and only a few as “intermediate”. Concerns about a similar change (breakpoint moved from 1 mg/L to 0.12 mg/L) in the Clinical and Laboratory Standards Institute (CLSI) interpretive thresholds have been expressed (38). There is a risk in over-categorizing isolates as penicillin-resistant, which would lead the clinician to use the broader spectrum macrolides. Moreover, the prescription of penicillin G appears appropriate to treat multi-microbial samples with *Streptococcus pyogenes*. According to the natural distribution of penicillin susceptibility values, the isolates from the present study should be considered as having a natural susceptibility phenotype (12, 38, 39).

Tetracycline resistance was observed in this study, concordant with other reports (13, 40, 12) but is clinically less relevant, given that recommended molecules such as amoxicillin were available. All isolates in which the *tetO* gene was found, were resistant to tetracycline. This gene encodes a protein that protects the ribosome from the action of tetracycline, allowing protein synthesis even in the presence of high intracellular tetracycline concentration (41).

The genotyping of isolates based on cgMLST provided confirmation or allowed to rule out suspected transmissions. In investigation group 1, the two isolates turned out to be phylogenetically very distant (as they belong to distinct sublineages), ruling-out transmission among the two cases. Instead of cross-transmission, the contact had asymptomatic carriage of a co-circulating *tox* negative strain. On the opposite, the isolates of investigation group 2 belonged to the same cgMLST genomic cluster (GC108). Although genomic clusters are defined using a tolerance of 25 mismatches, the isolates of this group were separated by only a maximum of 5 mismatches, underlying their very close genetic proximity.

Sublineage SL120, to which all five isolates in the *tox+* cluster (group 2) belong, was previously described only in two other isolates, one from 1995 in Finland and one isolated from 2015 in Australia; however, both belonged to other genomic clusters than GC499, which corresponds to group 2 isolates from New Caledonia. This illustrates the interest of classifying strains at both the sublineage and genomic group levels, as they provide information on genetic relationships, and hence transmission, at different timescales.

Our genomic analyses uncovered a high genetic diversity of *C. diphtheriae* circulating in New Caledonia, with at least 13 different sublineages, consistent with the situation in other localities (11–14). Nevertheless, nearly two thirds of the isolates belonged to only two predominant sublineages, SL228 and SL416. Besides, the latter had very restricted genetic diversity, as it comprised a single genomic cluster. The high frequency of these genotypes could be related to the insular nature of the New Caledonia territory. The fact that all SL416 isolates belong to the same genomic cluster suggests relatively recent transmission at the population scale (perhaps a few decades). Interestingly, SL416 corresponded to the isolates carrying the *tetO* gene, conferring tetracycline resistance; whether this phenotype has favored the spread of this group is an interesting possibility, as observed for other bacteria (42). SL416 was previously reported only once, isolated in France from a patient returning from French Polynesia.

Sublineage SL228 represented 21 isolates (36.2% of our samples) and has already been described elsewhere, sporadically. Of all the published data available worldwide (876 genomes), nine strains outside New Caledonia belong to this sublineage **(Table S3).** Their provenances are as follows: An isolate from Romania in 1966, two Australian isolates respectively from 2013 and 2016, two mainland France isolates from 2011 and 2021, an isolate from Wallis and Futuna from 2012, and three isolates from French Polynesia, isolated in 2017, 2019 and 2020. As observed for other pathogens, e.g., *Burkholderia pseudomallei* (43), the predominant sublineages seem to be linked either to geographically close territories (Polynesia, Australia) or from metropolitan France, consistent with human transmission and dissemination.

In this study, cases with *C. diphtheriae* had an uneven geographic repartition across New Caledonia. This observation can in part be explained by the geographical distribution of the population: high population density in the main city of Nouméa (34.7% of the population) and in the South Province (74.9 % of the population), but low density in the North Province and the Islands, with respectively 18.4 % and 6.8% of the population (15). *C. diphtheriae* is more easily detected with MALDI-TOF mass spectrometry, with which only the laboratories in Nouméa are equipped, including the laboratory that processed the samples from Lifou island. Whereas the Lifou clinicians regularly took samples from wounds with unfavorable evolution in order to seek bacteriological documentation, the other centers did so much less systematically. This bias can contribute to the overrepresentation of cases in Lifou, despite a small population (9,195 inhabitants in 2019, *i.e.,* 3.4% of the New Caledonian population) (15). This study therefore illustrates the difficulties of achieving a homogeneous diphtheria surveillance given existing heterogeneities of laboratory equipment, practices and awareness. Monitoring only symptomatic respiratory cases would have led to a very fragmentary picture, and the surveillance of diphtheria should clearly benefit from the addition of isolates from wound infections.

## Acknowledgements

We thank Simon Joxe for help with the geographical map, and Martine Noel (DASS-NC) and Yves Marie Ducrot (Province des iles) for help with epidemiological data completion. This work used the computational and storage services provided by the IT department at Institut Pasteur.

## Authors contributions

Conceptualization: ET, JC and SB; Methodology, ET, JC, EB, MH and SB; Genomic analyses, MH; Validation, JC and SB; Investigation, ET and MH.; Resources, JC, SB, JT and EB; Data Curation, ET; Writing – Original Draft Preparation, ET, MH, JC and SB; Writing – Review & Editing, all; Visualization, MH and ET; Supervision, JC and SB; Funding Acquisition, SB and JC.

## Author statement

This research was funded, in whole or in part, by Institut Pasteur and by European Union’s Horizon 2020 research and innovation programme. For the purpose of open access, the authors have applied a CC-BY public copyright license to any Author Manuscript version arising from this submission.

